# The congruency between anger intensity and reddish facial color modulates the early posterior negativity (EPN)

**DOI:** 10.64898/2026.06.15.732248

**Authors:** Rikuto Nishiura, Yuya Hasegawa, Hideki Tamura, Shigeki Nakauchi, Tetsuto Minami

**Author notes:** These authors contributed equally to this work (Rikuto Nishiura, Yuya Hasegawa). Conflicts of interest statement: The authors declare that they have no competing financial interests.

## Abstract

Facial color is associated with the perceptual evaluation of emotions, and compared with faces with original facial or greenish color, reddish angry faces are often judged as having higher emotion intensity. Although perceptual modulation by the relationship between anger and red has also been reported from the perspective of electroencephalography (EEG), how variations in perceived emotion intensity are reflected in brain activity remains unclear. This study investigated whether EEG activity associated with face and facial expression processing is modulated as a function of the interaction between facial color and perceived emotion intensity. In the experiment, we recorded EEGs while participants evaluated emotion intensity using facial stimuli created by combining morph continua from neutral to angry expressions with three types of facial color conditions (original, red, and green). The results revealed that, in the red facial color condition, the early posterior negativity (EPN) amplitude significantly increased as a function of emotion intensity compared with those in the original and green facial color conditions. These findings suggest that the early, automatic affective processing of facial expressions, reflected in the EPN, is modulated by the combination of facial color and emotion intensity. Our findings provide new evidence that early, automatic affective processing of facial expressions, as indexed by the EPN, is modulated by the congruency between high anger intensity and a reddish facial color.

**Highlight:**

- Reddish angry faces increase the ERP component associated with emotion evaluation.
- The relationship between anger and red is evident in the left hemisphere.
- The interaction between facial expression and color occurs at a later cognitive processing stage than facial expression or facial color processing alone.

## Introduction

Facial colors play important roles in social evaluations, facial expression recognition, and perceived emotion intensity (Nakajima et al., 2017; Benitez-Quiroz et al., 2018; Thorstenson and Pazda, 2021; Thorstenson et al., 2022; Shibusawa et al., 2025). In the judgment of angry faces, when the facial color is red, the face is more likely to be perceived as an angry face, even if the facial shape is the same (Nakajima et al., 2017; Peromaa and Olkkonen, 2019; Kato et al., 2022). Moreover, it has been reported that the perceived emotion intensity or aggressiveness of anger is increased by a reddish facial color (Thorstenson and Pazda, 2021; Thorstenson et al., 2021, 2022).

These effects of facial color are not confined to behavioral responses, as modulation of brain activity, such as the fusiform facial area and event-related potential (ERP) N170, depending on facial color, has also been reported (Nakajima et al., 2012, 2014). Recent research suggests that P300 amplitude related to selective attention is modulated by the interaction between facial expression and color (Hasegawa et al., 2025).

However, most previous studies have employed facial stimuli with neutral or categorically salient facial expressions. Therefore, differences in perceived emotion intensity, in other words, the strength of emotion inferred from facial expressions, remain unexamined. For example, although facial redness is known to increase the perceived emotion intensity of anger (e.g., Thorstenson et al., 2022), whether the brain wave activity underlying processes such as facial expression judgment and emotion recognition are modulated as a function of the relationship between facial color and emotion intensity remains unclear.

Among electroencephalogram (EEG) activities related to facial processing, several ERP components, including P100, N170, early posterior negativity (EPN) and P300, have been reported. P100 is a positive ERP amplitude observed approximately 100 ms after stimulus onset and is thought to reflect the processing of low-level visual features during the early stage of facial processing (Herrmann et al., 2005; Rossion and Caharel, 2011). N170 is associated with sensitivity to faces and is also modulated by facial expression and facial color, with greater amplitudes for emotional expressions such as angry faces than for neutral faces (Bentin et al., 1996; Hinojosa et al., 2015; Schindler and Bublatzky, 2020). The EPN has a negative amplitude, which is related to early selective processing of emotional stimuli and is enhanced by emotional faces (Schupp et al., 2004; Foti et al., 2009; Langeslag et al., 2018). P300 is a positive amplitude associated with top-down processes such as selective attention. P300 has also been linked to facial expression processing, with increased amplitudes observed for angry or fearful expressions (Rossignol et al., 2005; Chai et al., 2012; Lin et al., 2020). Although these ERP components associated with face and facial expression processing have been reported, whether they are modulated by the relationship between the perceived emotion intensity of facial expressions and facial color remains unclear.

Hence, this study used morphed facial expression stimuli and recorded EEGs during a task involving the evaluation of the perceived emotion intensity of the stimuli. It has been reported that perceived responses to angry faces are enhanced by increased facial redness (Nakajima et al., 2017; Thorstenson and Pazda, 2021; Thorstenson et al., 2022; Hasegawa et al., 2025; Shibusawa et al., 2025). Accordingly, focusing on the interaction between anger and red, we hypothesized that the increase in the perceived emotion intensity of anger by reddish facial color would be reflected in ERP components associated with face and facial expression processing, including N170, EPN, and P300. In particular, we hypothesized that high perceived emotion intensity would be predicted from ERP components in response to the combination of high emotional anger intensity and reddish facial color. This expectation was based on the assumption that this increase, driven by the enhanced salience of anger representations manipulated using morphed stimuli, would be further amplified by reddish facial color.

## Method

### Participants

Twenty Japanese students (1 female, 19 male, mean age = 22.4 ± 1.3 years) at Toyohashi University of Technology participated in the experiment. The sample size was set based on a previous study (Hasegawa et al., 2025). This experiment was conducted with the approval of the Ethics Review Committee for Research Involving Human Subjects at Toyohashi University of Technology.

### Stimuli

The facial stimuli are illustrated in Figure 1. Neutral and angry facial images of two Japanese individuals (1 female and 1 male) from the ATR Facial Expression Image Database (ATR-Promotions; Kyoto, Japan; https://www.atr-p.com/products/face-db.html) were used. As facial expression stimuli, the facial images were morphed from neutral to angry in 3 steps by using MATLAB 2021a. Before morphing, we trimmed the facial images into oval shapes by using Photoshop (Adobe Inc., San Jose, CA, USA) to remove the hair, ears, and necks. The average image luminance was controlled via SHINE_color, a MATLAB toolbox (Willenbockel et al., 2010; Ben, 2021). Based on an experiment by Hasegawa et al. (2025), we manipulated the facial color of the stimuli across three facial color conditions: original (no manipulation, CIELAB a*±0 unit), red (a*+12 unit), and green (a*−12 unit). Therefore, a total of 18 facial images (3 angry levels × 3 facial colors × 2 individuals) were presented as experimental stimuli. The image dimensions were 4.2° × 5.5°, and the background color was always gray (*Y* = 17.09 cd/*m*^2^).

**Figure 1.**
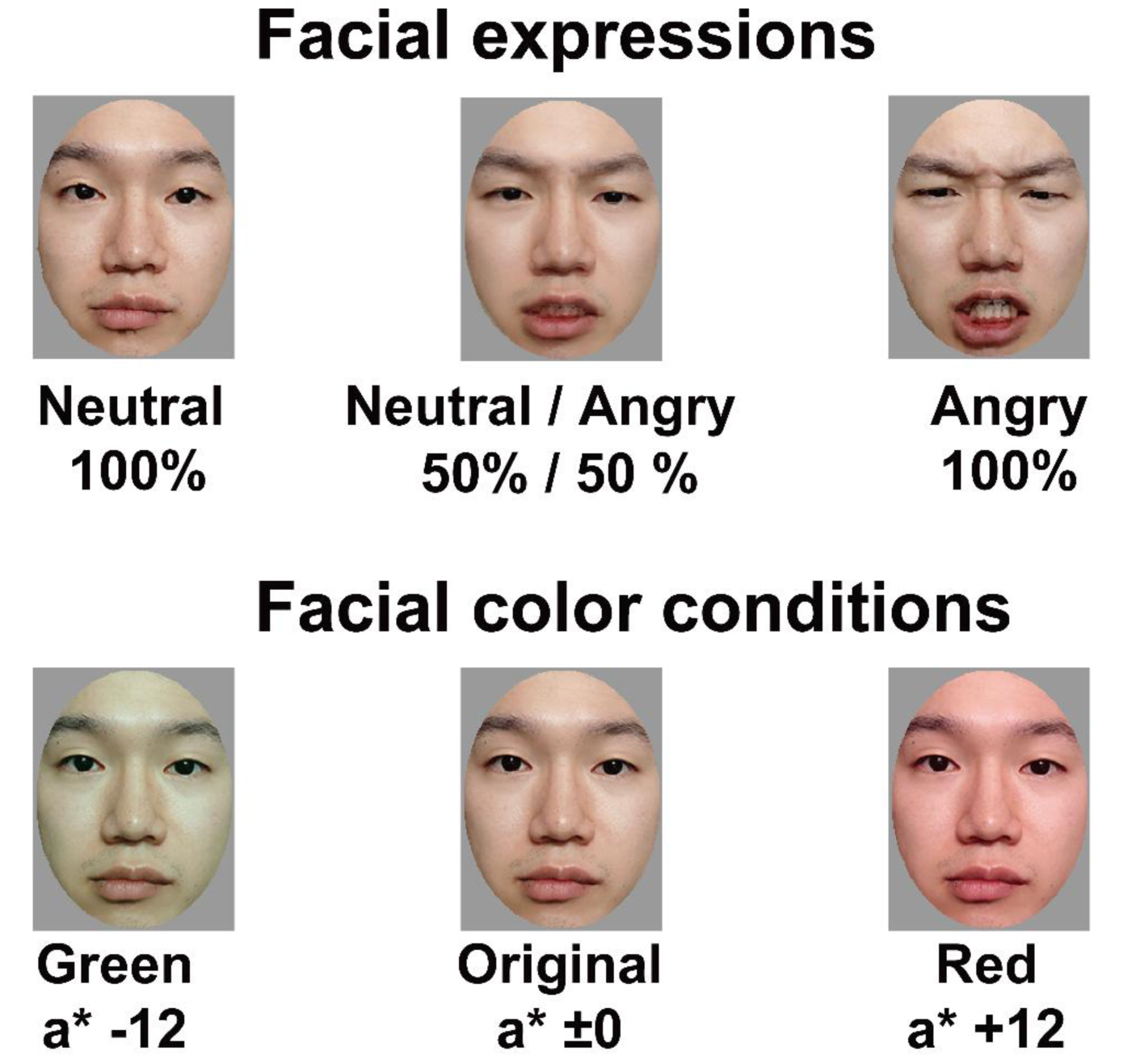
Examples of the image stimuli for each condition. A 3-step (neutral to angry) morphing of facial stimuli was used. Four facial color conditions were provided using green, the original color, and red. The face in the figure is from one of the authors (Y.H.) with his permission and was not used in the experiment.

### Apparatus

A dark, magnetically shielded room was used in the experiment. The stimuli were presented on a monitor (VIEPixx/EEG, VPixx Technologies Inc., Canada; resolution: 1920 × 1080 ; frame rate 120 Hz; white point [*x*, *y*] = [0.30, 0.33], *Y* = 91.23 cd/m^2^). Participants completed the task in a seated position with their heads stabilized using a chin rest placed 60 cm from the monitor. Stimulus presentation and experimental control were implemented using Psychtoolbox 3.0.17 (Brainard, 1997; Pelli, 1997; Kleiner et al., 2007). EEG signals were recorded using a BioSemi ActiveTwo system (BioSemi, Amsterdam, Netherlands) with 64 channels of electrodes and 6 channels of external sensors, sampled at a rate of 512 Hz.

### Procedure

In the experiment, participants rated the emotion intensity of each facial stimulus on a seven-point scale (“1” is low, “7” is high). A summary of the experimental procedure is shown in Figure 2. Each trial began with a 1.0-second interstimulus interval (ISI), followed by a 1.0-second presentation of a fixation and a 2.0-second presentation of facial stimulus. After the facial stimulus was presented, the participants responded to the emotion intensity of the presented stimulus using a numeric keypad. Each facial stimulus was rated two times per participant. In total, the participants performed 36 trials of the evaluation task. EEG data were recorded during the task.

**Figure 2.**
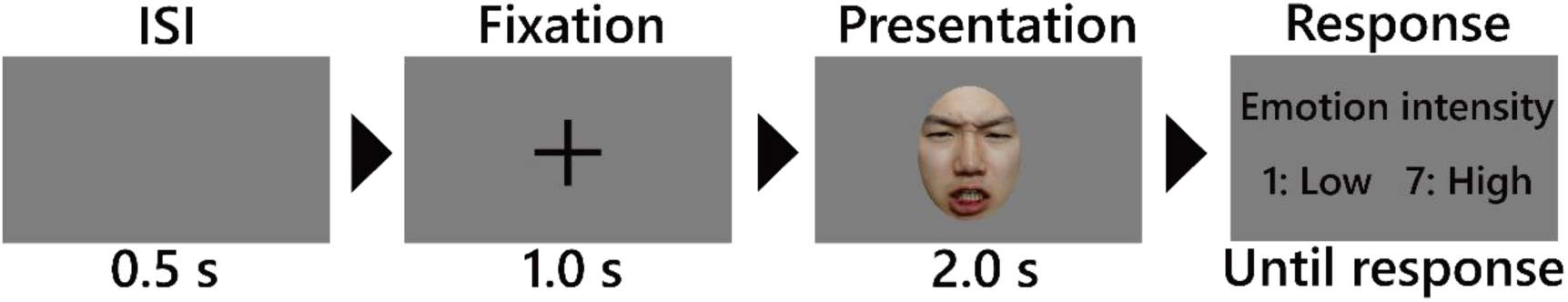
Experimental Procedure. Notably, the ratio of the screen to the stimulus depicted in this figure differs from the actual ratio. The face in the figure is from one of the authors (Y.H.) with his permission and was not used in the experiment.

### Preprocessing of EEG data

For preprocessing, we downsampled the EEG data to 200 Hz. Afterward, a high-pass filter with a cutoff frequency of 1 Hz was applied, and the “cleanLineNoise” function in EEGLAB was used to attenuate line-related interference, including broadband white noise, power line noise at 60 Hz, and its harmonic components (120, 180, and 240 Hz), with the significance cutoff level being *p* = .01. Electrodes exhibiting poor signal quality were identified and excluded using the “clean_rawdata” function in EEGLAB. The criteria applied included a flatline duration threshold of 5 s, a minimum correlation of 0.85 with surrounding electrodes, a deviation exceeding four standard deviations from the mean, and artifact rejection based on the ASR algorithm. Channels rejected by “clean_rawdata” were subsequently reconstructed using spherical spline interpolation based on information from surrounding electrodes. Moreover, artifacts were further reduced by identifying and removing ocular components through adaptive mixture independent component analysis (AMICA) in combination with ICLabel (Leutheuser et al., 2013; Pion-Tonachini et al., 2019). Finally, we set the time of stimulus presentation as 0 ms and extracted the EEG data at −100∼1000 ms. Finally, stimulus onset was defined as 0 ms , and EEG epochs were extracted from −100∼1000 ms relative to stimulus presentation.

One participant was excluded from further analysis because over 50% of their EEG data were removed during preprocessing. For the remaining participants, the mean proportion of missing EEG data was 6.14%.

## ERP Extraction

### P100

First, channel-averaged EEG signals were extracted from Iz, Oz, O1, O2, and POz based on the preprocessed data. The baseline EEG was defined as the average EEG of −100∼0 ms. The peak amplitudes were then computed for each trial within the 90∼140 ms window following the onset of facial stimuli. The selection of time windows and electrode sites was determined a priori during the experimental design phase, drawing on findings from prior studies (Polich, 2007; Hinojosa et al., 2015; Che et al., 2024; Hasegawa et al., 2025).

### N170 (Left, Right)

The channel-averaged EEG at 5 channels near the temporal area (Left: TP7, P5, P7, P9, PO7; Right: TP8, P6 P8, P10, PO8) was extracted from the preprocessed EEG. The peak amplitudes at 150∼200 ms after the presentation of the target stimuli were then calculated for each trial. The other parameters were the same as those of P100.

### EPN (Left, Right)

We extracted the channel-averaged EEG at 2 channels near the temporal area (Left: P7, PO7; Right: P8, PO8) from the preprocessed EEG. The baseline time window was the same as that for the P100 and N170 amplitudes. The peak amplitudes at 250∼300 ms after the facial stimuli were presented were then calculated for each trial.

### P300

Based on a previous study, we extracted the channel-averaged EEG at Cz, CPz, CP1, CP2, and Pz from the preprocessed EEG and averaged the amplitudes at 300∼500 ms after the presentation of facial stimuli for each trial (Hasegawa et al., 2025). The others (e.g., baseline) were the same as the other ERPs.

#### Statistical analysis

First, the rating of emotion intensity was predicted based on ERP and facial color. We applied a linear mixed-effects model (LMM) to each rating within the three facial colors (original, red, green), the six ERPs (P100, N170 Left, N170 Right, EPN Left, EPN Right, P300), and all participants. The model included facial color, ERPs, and all interaction terms between facial color and each ERP (not between ERPs) as fixed effects, along with a random intercept for participants. T tests were performed using Satterthwaite’s method. The model was constructed as follows:

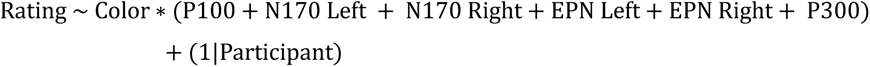

After the ERP components and facial color conditions associated with ratings were assessed using the predictive model, we further analyzed ERP modulations in detail by predicting each ERP component. We applied an LMM to single-trial ERP amplitudes within the ratings of perceived emotion intensity, facial colors, and all participants, with independent analyses performed for each ERP component. The model included the rating (rescaled per participant), facial color, and their interaction as fixed effects and a random intercept for participants. We considered that the effects of facial shape (morphing about facial expression) and individual facial stimuli are included in the rating of emotion intensity. The model was constructed as follows:

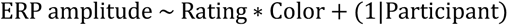

A Type III analysis of variance with Satterthwaite’s method was subsequently conducted for the calculated model. All the statistical tests were two-tailed, the significance threshold was set at 0.05, and the Holm method was used as the p value adjustment method for the post hoc test. LMM analyses were conducted in R using the lmerTest, lme4 and emmeans packages.

## Results

### Rating

Before the LMM analysis, we plotted emotion intensity ratings at each morphing level to confirm that the facial expression morphing stimuli functioned as intended (see Figure 3). As expected, the ratings of emotion intensity increased with increasing levels of anger, and higher ratings were observed in the red facial color condition than in the original color and green conditions.

**Figure 3.**
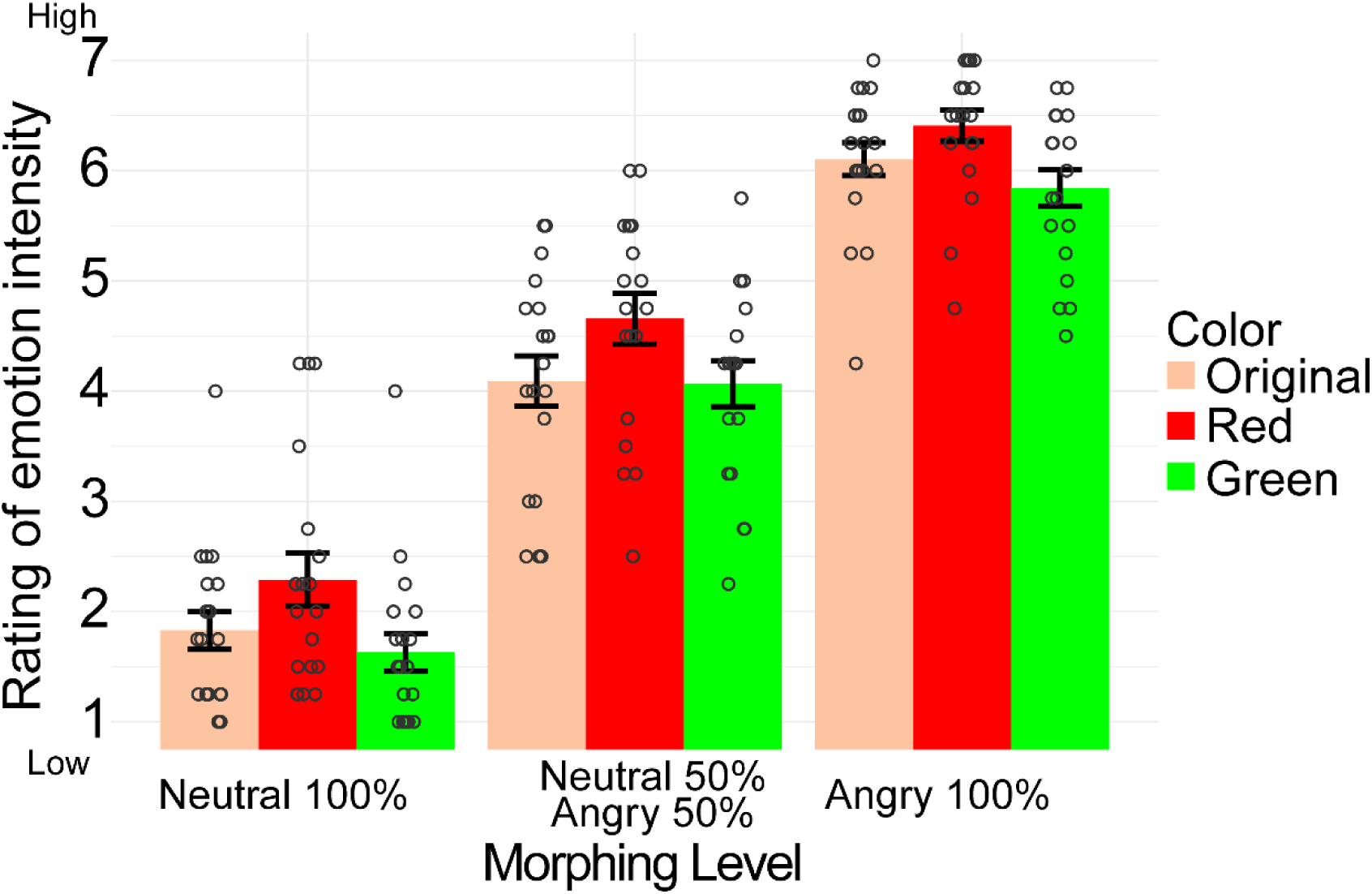
Participants’ rating means for each facial stimulus. The points of each gray circle represent individual data, and error bars show the standard error of the mean.

The results of the LMM for ratings revealed a significant interaction between facial color and EPN Left (original vs. red: *t*(314.2) = −2.313, *Pr*(> |*t*|) = .021, *Estimate* = −0.137 ). The results suggest that the relationship between the left-hemisphere EPN and ratings of perceived emotion intensity is modulated by facial color, particularly with reddish facial color enhancing the EPNs’ negative amplitude. The other results and the results of the generalized variance inflation factor (GVIF) are shown in Tables 1–2.

**Table 1.** Fixed effects of the fitted linear mixed model (LMM)

|  | Estimate | Std.error | df | t value | Pr (>\|t\|) |
| --- | --- | --- | --- | --- | --- |
| Intercept | 3.563 | 0.513 | 269.07 | 6.951 | $2.73 \times 10^{-11}$ |
| Red | 0.867 | 0.739 | 309.43 | 1.174 | 0.241 |
| Green | 0.276 | 0.716 | 313.82 | 0.385 | 0.701 |
| P100 | 0.023 | 0.045 | 315.29 | 0.523 | 0.602 |
| N170_Left | -0.100 | 0.044 | 319.00 | -2.177 | 0.030 |
| N170_Right | 0.011 | 0.038 | 305.04 | 0.301 | 0.764 |
| EPN_Left | 0.094 | 0.044 | 317.24 | 2.107 | 0.036 |
| EPN_Right | -0.039 | 0.036 | 318.01 | -1.092 | 0.276 |
| P300 | 0.054 | 0.069 | 310.50 | 0.771 | 0.441 |
| Red:P100 | -0.029 | 0.063 | 315.13 | -0.466 | 0.641 |
| Green:P100 | -0.015 | 0.059 | 317.78 | -0.256 | 0.799 |
| Red:N170_Left | 0.091 | 0.063 | 316.15 | 1.454 | 0.147 |
| Green:N170_Left | 0.097 | 0.064 | 317.47 | 1.517 | 0.130 |
| Red:N170_Right | -0.019 | 0.061 | 315.64 | -0.310 | 0.757 |
| Green:N170_Right | -0.042 | 0.060 | 312.57 | -0.709 | 0.479 |
| Red:EPN_Left | -0.133 | 0.060 | 311.79 | -2.207 | 0.028 |
| Green:EPN_Left | -0.017 | 0.066 | 315.45 | -0.254 | 0.799 |
| Red:EPN_Right | 0.041 | 0.052 | 313.60 | 0.790 | 0.430 |
| Green:EPN_Right | 0.017 | 0.056 | 317.08 | 0.307 | 0.759 |
| Red:P300 | 0.095 | 0.095 | 318.99 | 1.001 | 0.318 |
| Green:P300 | -0.080 | 0.105 | 319.00 | -0.766 | 0.444 |

**Table 2.** Results of the generalized variance inflation factor (GVIF)

| | GVIF | Df | $GVIF^{1/(2 \cdot Df)}$ |
| --- | --- | --- | --- |
| Color | 54.29 | 2 | 2.71 |
| P100 | 4.19 | 1 | 2.05 |
| N170_Left | 4.00 | 1 | 2.00 |
| N170_Right | 2.96 | 1 | 1.72 |
| EPN_Left | 5.06 | 1 | 2.25 |
| EPN_Right | 3.63 | 1 | 1.90 |
| P300 | 3.23 | 1 | 1.80 |
| Color:P100 | 26.92 | 2 | 2.28 |
| Color:N170_Left | 15.98 | 2 | 2.00 |
| Color:N170_Right | 15.93 | 2 | 2.00 |
| Color:EPN_Left | 9.64 | 2 | 1.76 |
| Color:EPN_Right | 5.69 | 2 | 1.54 |
| Color:P300 | 4.13 | 2 | 1.43 |

All adjusted GVIF values were less than 5 (max is Color: 2.71), suggesting that multicollinearity was not a major concern.

### P100

The model results for the relationship between ERP P100 and the rating of emotion intensity are shown in Figure 4. The x-axis ranged from −4– to 4 because ratings were mean centered within participants prior to the LMM analysis. This procedure was applied to standardize the central tendency of judgments and reduce interindividual variability in rating tendencies. Across participants, the mean ratings ranged from 3.12–5.00. No significant main effect or interaction effect of facial expression and facial color was detected (Color: *F*(2, 316.14) = 0.112, *p* = .894, *η*_*p*_^2^ = 7.10 × 10^−4^; Rating: *F*(1, 316.06) = 0.034, *p* = .853, *η*_*p*_^2^ = 1.08 × 10^−4^; Interaction: *F*(2, 318.95) = 0.616, *p* = .539, *η*_*p*_^2^ = 3.86 × 10^−3^).

**Figure 4.**
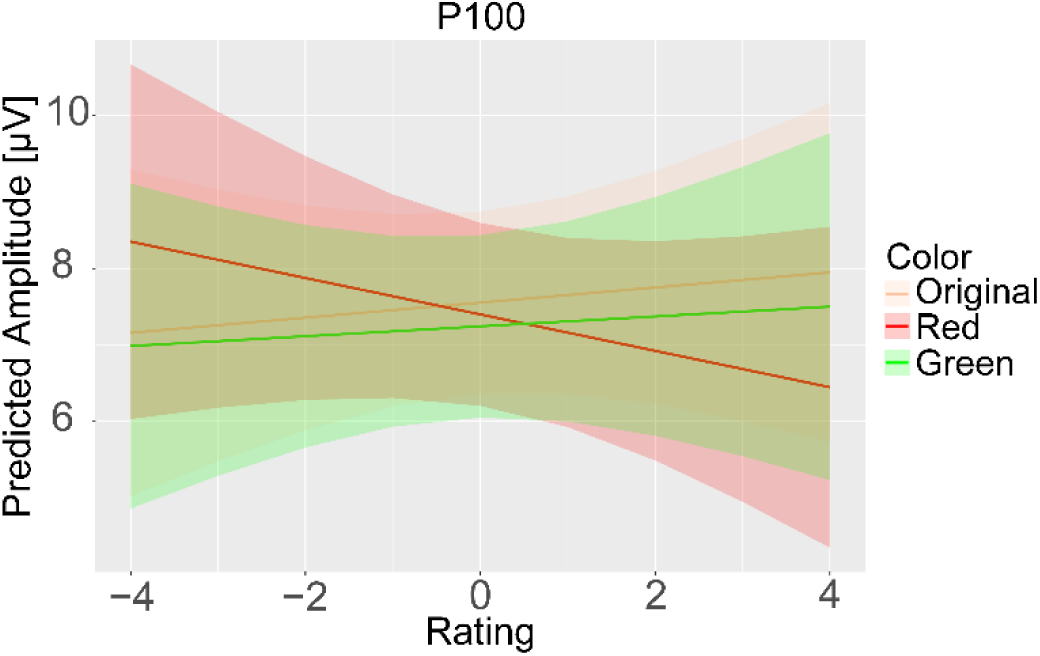
The predicted model results of P100. The vertical axis represents the predicted P100 amplitude [μV], and the horizontal axis represents emotion intensity ratings (centered at 0). The lines represent model-based marginal predictions, with the shaded bands indicating the corresponding confidence intervals.

### N170

The model results for the relationship between N170s and the rating of emotion intensity are shown in Figure 5. On the left-side N170, no significant main effect or interaction effect of facial expression and facial color was observed (Color: *F*(2, 316.09) = 0.886, *p* = .413, *η*_*p*_^2^ = 5.58 × 10^−3^; Rating: *F*(1, 316.04) = 1.563, *p* = .212, *η*_*p*_^2^ = 4.92 × 10^−3^; Interaction: *F*(2, 317.67) = 1.825, *p* = .163, *η*_*p*_^2^ = 0.01). In addition, there were no significant main effects or interaction effects of facial expression on the left-side N170 (Color: *F*(2, 316.05) = 1.191, *p* = .305, *η*_*p*_^2^ = 7.48 × 10^−3^; Rating: *F*(1, 316.02) = 2.493, *p* = .115, *η*_*p*_^2^ = 7.83 × 10^−3^; Interaction: *F*(2, 317.21) = 0.141, *p* = .868, *η*_*p*_^2^ = 8.89 × 10^−4^).

**Figure 5.**
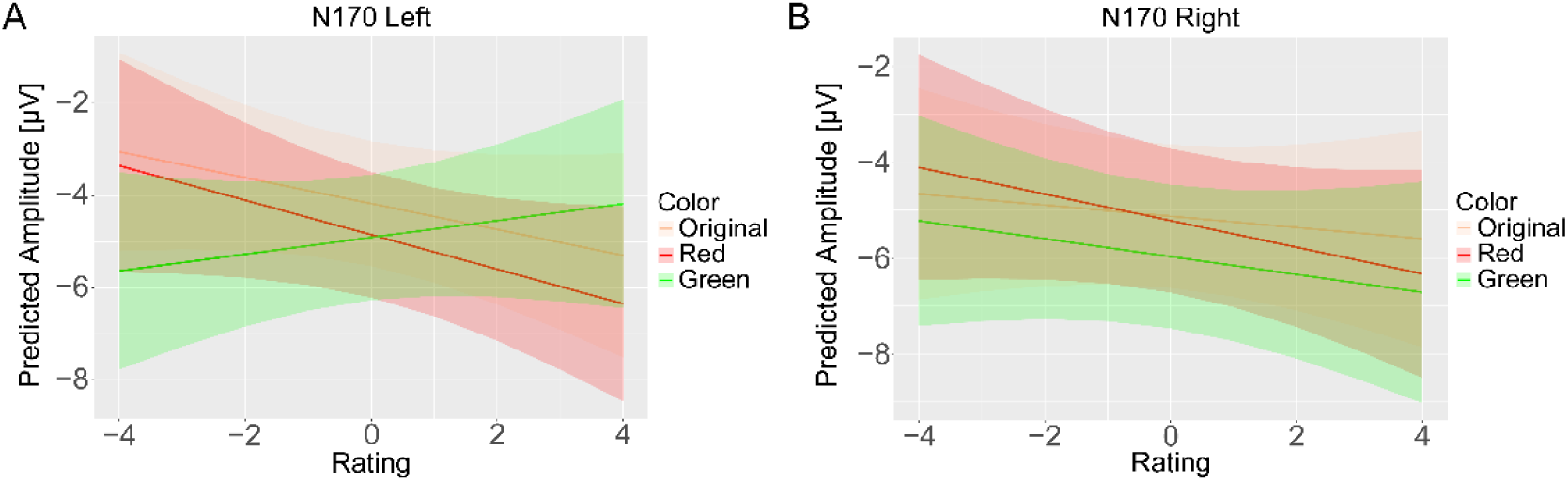
The predicted model results of N170 (A: Left, B: Right). The vertical axis represents the predicted N170 amplitude [μV]. The others were the same as in Figure 2.

### EPN

The model results of the relationship between the EPN and the rating of emotion intensity are shown in Figure 6. We found no significant main effect of facial color ( *F*(2, 316.11) = 0.991, *p* = .372, *η*_*p*_^2^ = .006) or rating (*F*(1, 316.03) = 0.701, *p* = .403, *η*_*p*_^2^ = .002). However, there was a significant interaction effect between the rating value and facial color (*F*(2, 319.11) = 4.255, *p* = .015, *η*_*p*_^2^ = .026). The post hoc results revealed that the increase in the left-side EPN (i.e., increased negative amplitude) associated with increasing perceived emotion intensity was significantly greater when the facial color was red than when it was the original color or green (red vs. original: *t*(319) = 2.472, *p* = .031, Cohen^′^s *d* = 0.167 ; red vs. green: *t*(319) = 2.586, *p* = .031, Cohen^′^s *d* = 0.175). There was no significant difference between the original and green facial color conditions (*t*(319) = 0.131, *p* = .896, Cohen^′^s *d* = 0.009).

**Figure 6.**
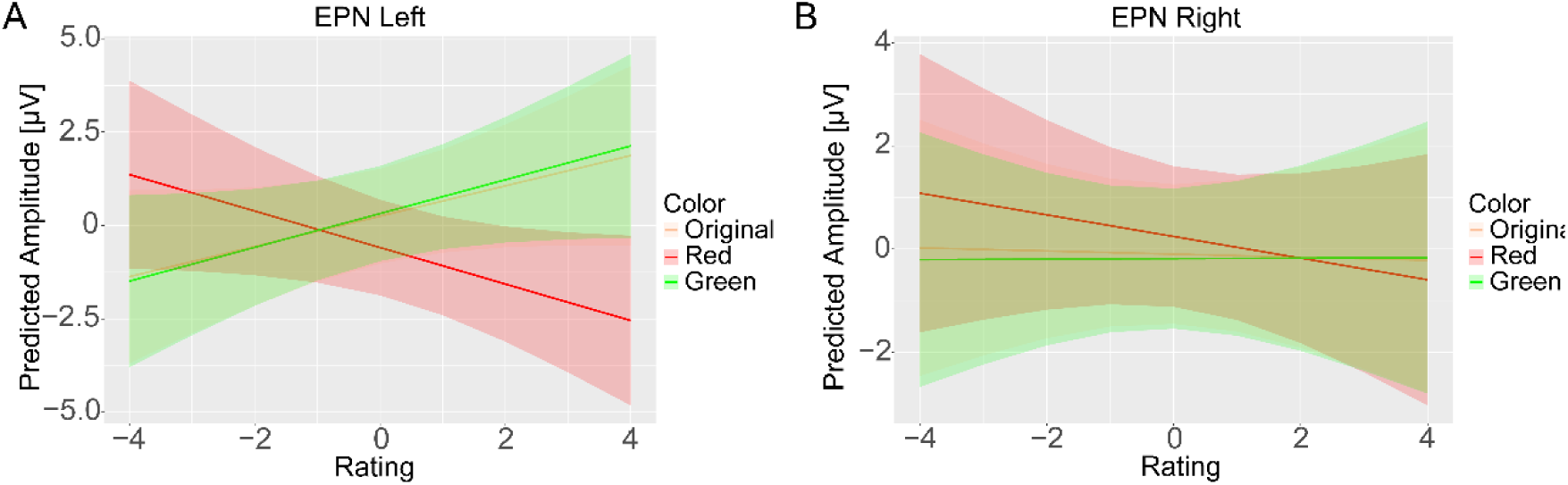
The predicted model results of the EPN (A: Left, B: Right). The vertical axis represents the predicted EPN amplitude [μV]. The others were the same as in Figure 2.

In contrast, no main effects or significant interactions were detected on the right-side EPN (Color: *F*(2, 315.98) = 0.172, *p* = .842, *η*_*p*_^2^ = .001 ; Rating: *F*(1, 315.89) = 0.246, *p* = .620, *η*_*p*_^2^ = 7.77 × 10^−4^; Interaction: *F*(2, 319.09) =, *p* = 0.174, *η*_*p*_^2^ = .001).

### P300

The model results for the relationship between P300 and the rating of emotion intensity are shown in Figure 7. We found a significant main effect of the emotion intensity rating (*F*(1, 315.93) = 4.407, *p* = .037, *η*_*p*_^2^ = 0.01), and the results revealed that increasing perceived emotion intensity led to a greater P3 amplitude. However, no main effect of color (*F*(2, 316.04) = 2.763, *p* = .065, *η*_*p*_^2^ = 0.02) and no significant interaction between color and rating were observed (*F*(2, 315.79) = 1.321, *p* = .268, *η*_*p*_^2^ = 8.20 × 10^−3^).

**Figure 7.**
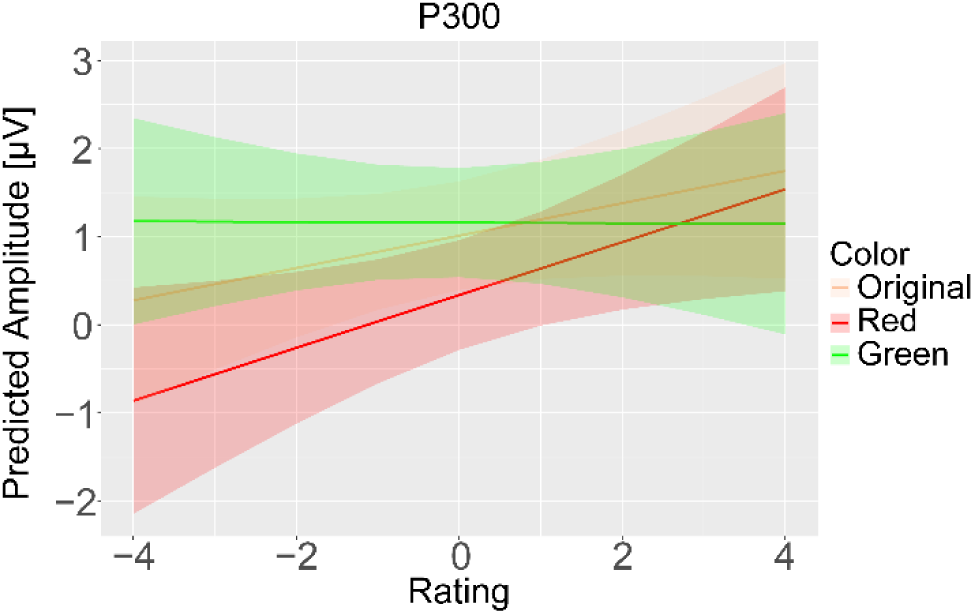
The predicted model results of P300. The vertical axis represents the predicted P300 amplitude [μV]. The others were the same as in Figure 2.

## Discussion

In this study, we recorded EEGs during an emotion intensity evaluation task using morphed facial expression stimuli with different facial colors to investigate ERP modulations as a function of increases in the perceived emotion intensity of anger driven by the interaction between anger and red. On the EPN, we found a significant interaction between emotion intensity and color, and the EPN amplitude increased as emotion intensity increased when the facial color was reddish. These findings might be attributable to contributions from the processing of angry faces, such as emotion processing and early automatic attention. The EPN is associated with emotional evaluation, and larger EPN amplitudes have been reported in response to emotional facial expressions, such as angry and happy faces, as well as to the valence and affective meaning of stimuli (Schupp et al., 2004; Foti et al., 2009; Palazova et al., 2011). In addition, it has been reported that the EPN differed depending on the facial features of angry faces, such as whether the mouth was open or closed (Langeslag et al., 2018).

Therefore, facial expression and facial color may have served as cues for emotion evaluation, and the EPN may have reflected local processing of these facial features and increasing perceived emotion intensity driven by reddish angry faces. It has been suggested that facial color influences emotion perception as a factor independent of facial muscle movements (Benitez-Quiroz et al., 2018). Taken together, the increase in the EPN observed in our study might be attributable to the fact that the processing of local facial features, such as facial expression and facial color, during emotion intensity evaluation was more likely to match participants’ affective meaning of angry faces with high emotion intensity.

In terms of the relationship between angry and facial color, from a physiological perspective, facial color is related to blood flow, and increased facial redness reflects increased blood flow due to emotional arousal (Edwards and Duntley, 1939; Drummond et al., 2001). Furthermore, from a psychological perspective, numerous previous studies have reported that facial redness enhances perceptual responses (e.g., facial expression recognition, emotion intensity, and social evaluations) to angry faces (Nakajima et al., 2017; Thorstenson and Pazda, 2021; Thorstenson et al., 2022; Hasegawa et al., 2025). In addition, angry faces may be remembered as having a more reddish-yellowish facial color than neutral faces do (Hasegawa et al., 2024). Hence, our findings suggest that facial color modulates brain activity associated with emotion processing and affective meaning. In particular, our results provide neural findings supporting the relationship between facial color and emotion intensity, especially the enhancement of anger perception induced by facial redness.

In the EPN results, a significant interaction between emotion intensity and facial color was observed in the left hemisphere, whereas no significant main effects or interactions were detected in the right hemisphere. One possible factor that influence these results is the asymmetry of interhemispheric function. Although the EPN has often been reported as bilateral or right-lateralized for emotional stimuli (Schupp et al., 2004), feature-based and semantic processing of faces has also been associated with the left hemisphere (Jiang et al., 2017). The left-lateralized interaction we observed should therefore be considered a tentative observation that requires direct hemisphere-by-color statistical testing in future work. The manipulation of facial color in our experiment can be regarded as feature information added while maintaining the original facial structure (i.e., facial shape). Thus, it is reasonable to assume that the interaction between facial expression and facial color was more strongly reflected in the left hemisphere, which is associated with the processing of local features and semantic integration, than in the right hemisphere, which is associated with the processing of the global structure of faces.

Our results showed that P300 amplitudes increased with increasing perceived emotion intensity. These results were similar to those of previous studies that reported an increase in P300 amplitudes in response to angry faces (Chai et al., 2012; Lin et al., 2020; Hasegawa et al., 2025). However, there was no interaction between anger (high emotion intensity) and red, as observed by Hasegawa et al. (2025). Hasegawa et al. (2025) employed an oddball paradigm, which is commonly used to elicit the P300 component. Thus, their experimental task consisted of category comparisons under sequential dependence, in which the updating of working memory or selective attention played a central role.

Conversely, our experimental task involved the evaluation of perceived emotion intensity in facial stimuli and was related to the contextual processing of facial expressions. The EPN reflects early, automatic attentional enhancement to emotionally salient stimuli (Schupp et al., 2004; Schindler and Bublatzky, 2020), whereas the P300 index later reflects top-down processes such as selective attention and working memory updating in facial categorization (Polich, 2007; Kaufmann et al., 2009). Moreover, it has also been suggested that face-related ERPs are modulated by task context (Schindler and Bublatzky, 2020). Therefore, the modulation of brain activity by the interaction between anger and red depends on the task context.

Our findings revealed that the EPN amplitude increased in response to the reddish facial color and high emotion intensity of anger. In contrast, these effects were not observed in earlier ERP components (P100 and N170). These results were similar to those of a previous study and suggested that the interaction between anger and red contributes to the later cognitive processing stage than the processing of facial expression or facial color alone does (Hasegawa et al., 2025). In particular, P100 and N170 are primarily associated with early bottom-up perceptual processing of structural facial information, whereas the EPN reflects an automatic, attention-related enhancement to emotionally salient stimuli that emerges later in the visual processing stream (Schupp et al., 2004; Schindler and Bublatzky, 2020). Therefore, the interaction between anger and red in the EPN window suggests that the affective salience signal indexed by the EPN is amplified when expression and color combinations are perceptually coherent rather than requiring direct top-down access to memory representations.

The limitations of this study are as follows. First, the experiments in this study were restricted to the combination of anger and red, to which a salient response was elicited in previous studies (e.g., Nakajima et al., 2017; Hasegawa et al., 2025). However, the relationship between emotion and color is not limited to anger and red. Combinations such as happy expressions with yellow (or red) and sadness with blue have also been reported (Thorstenson et al., 2018). Previous studies have reported that facial colors congruent with specific expressions enhance the accuracy of emotion categorization while increasing perceived emotion intensity and influencing social evaluations (Thorstenson and Pazda, 2021; Thorstenson et al., 2021, 2022; Nguyen et al., 2023). Therefore, the suggestion of this study should be limited to the finding that the combination of anger and red modulated the EPN. Further investigation is needed before it can be concluded that brain activity is modulated by the congruency between facial expression and color.

Second, our results do not allow us to clearly determine whether the observed modulations were driven by physiological or semantic congruency. Because our experiment was based on the experiment of Hasegawa et al. (2025), it does not consider natural changes in facial color. Changes in facial color in nature are not uniform across the face (MORETTI et al., 1959; Jimenez et al., 2010; Kikuchi et al., 2015). It has been reported that facial blood flow increases during anger expression, resulting in increased facial redness (Drummond, 1999; Drummond et al., 2001; Kreibig, 2010). However, such color change is determined not only by a single hue (i.e., not only a* channel) but also by factors such as hemoglobin in the blood flow and melanin in the skin (Edwards and Duntley, 1939; Zonios et al., 2001; Kikuchi et al., 2015). Moreover, facial color changes are not uniform due to variations in the thickness of facial skin and the distribution of blood vessels (MORETTI et al., 1959; Tsumura et al., 1999, 2003). Therefore, it should be noted that our findings do not necessarily reflect congruency with physiological factors.

Third, our sample was highly imbalanced in terms of sex (19 male participants and 1 female participant). Because sex differences in facial processing, color perception, and emotion judgment have been reported, our findings cannot be generalized beyond a predominantly male population, and sex effects cannot be tested with the present data. Future studies should recruit sex-balanced samples to evaluate the robustness of the observed interaction between anger and red.

Fourth, the individual characteristics of the participants were not assessed in this experiment. The ability to detect faces, as well as to recognize and process facial expressions and to show attentional biases toward facial expressions such as angry faces, varies depending on factors such as trait anxiety, Moebius syndrome, or autism spectrum disorder (Golarai et al., 2006; Surcinelli et al., 2006; Telzer et al., 2008; Tanaka and Sung, 2016; Quettier et al., 2023). Therefore, the individual characteristics of the participants may have influenced the processing of facial recognition or memory referral, and the extent to which these characteristics affect the magnitude of the observed effects should be examined in future studies. These individual characteristics were not directly measured in the present study and could not be statistically controlled for; the random intercepts in our model account only for overall amplitude offsets per participant and do not absorb structured between-subject variation related to such traits.

## Conclusion

We investigated how brain activity is modulated as a function of the relationship between levels of emotion intensity and facial color, particularly between anger and red. The results suggest that the EPN, an ERP index of automatic affective processing, is modulated by the combination of high anger intensity and reddish facial color.

### Commercial relationships

None.

### Author contributions

Y.H. and T.M. designed the research; R.N. and Y.H. provided experimental code and conducted the analysis; R.N., Y.H. and T.M. wrote the paper; H.T. and S.N. provided feedback on the manuscript.

Y.H., H.T., S.N. and T.M. were responsible for funding acquisition.

### Data availability

The analysis code and the data used for statistical analysis are available at https://osf.io/t53uq/. However, the data are publicly available only as amplitude extraction data for each experimental condition of each participant, in accordance with the regulations of the Ethics Committee for Human Research at Toyohashi University of Technology.

## Acknowledgment

This work was supported by JSPS KAKENHI (Grant Numbers JP26KJ1371 to Y.H., JP25K21323 to H.T., JP25H01141 to S.N., JP26H00512 to T.M., and JP23KK0183 to T.M.), JST SPRING, Japan (Grant Numbers JPMJSP2171 to Y.H.), and the student fellowship program for the Leading Graduate School at Toyohashi University of Technology to Y.H.

## Notes

### Competing Interest Statement

The authors have declared no competing interest.

## References

Ben RD (2021) SHINE _ color : controlling low-level properties of colorful images.

Benitez-Quiroz CF, Srinivasan R, Martinez AM (2018) Facial color is an efficient mechanism to visually transmit emotion. Proc Natl Acad Sci U S A 115:3581–3586.

Brainard DH (1997) The Psychophysics Toolbox. Spat Vis 10:433–436.

Chai H, Chen WZ, Zhu J, Xu Y, Lou L, Yang T, He W, Wang W (2012) Processing of facial expressions of emotions in healthy volunteers: An exploration with event-related potentials and personality traits. Neurophysiologie Clinique/Clinical Neurophysiology 42:369–375.

Che J, Cheng N, Jiang B, Liu Y, Liu Haihong, Li Y, Liu Haining (2024) Executive function measures of participants with mild cognitive impairment: Systematic review and meta-analysis of event-related potential studies. International Journal of Psychophysiology 197:112295.

Drummond PD (1999) Facial flushing during provocation in women. Psychophysiology 36:325–332.

Drummond PD, Quah SH, Drummond P (2001) The effect of expressing anger on cardiovascular reactivity and facial blood flow in Chinese and Caucasians. Psychophysiology 38:190–196.

Edwards EA, Duntley SQ (1939) The pigments and color of living human skin. American Journal of Anatomy 65:1–33.

Foti D, Hajcak G, Dien J (2009) Differentiating neural responses to emotional pictures: Evidence from temporal-spatial PCA. Psychophysiology 46:521–530.

Golarai G, Grill-Spector K, Reiss AL (2006) Autism and the development of face processing. Clin Neurosci Res 6:145–160.

Hasegawa Y, Tamura H, Nakauchi S, Minami T (2025) Interaction between Facial Expression and Color in Modulating ERP P3. eNeuro 12:ENEURO.0419-24.2024.

Hasegawa Y, Tamura H, Nakauchi S, Minami T (2024) Facial expressions affect the memory of facial colors. J Vis 24:14–14.

Herrmann MJ, Ehlis AC, Muehlberger A, Fallgatter AJ (2005) Source localization of early stages of face processing. Brain Topogr 18:77–85.

Hinojosa JA, Mercado F, Carretié L (2015) N170 sensitivity to facial expression: A meta-analysis. Neurosci Biobehav Rev 55:498–509.

Jiang L, Wang Yun, Cai B, Wang Yueming, Wang Yiwen (2017) Spatial-temporal feature analysis on single-trial event related potential for rapid face identification. Front Comput Neurosci 11:290696.

Jimenez J, Scully T, Barbosa N, Donner C, Alvarez X, Vieira T, Matts P, Orvalho V, Gutierrez D, Weyrich T (2010) A practical appearance model for dynamic facial color. ACM Trans Graph 29.

Kato M, Sato H, Mizokami Y (2022) Effect of skin colors due to hemoglobin or melanin modulation on facial expression recognition. Vision Res 196:108048.

Kaufmann JM, Schweinberger SR, Burton AM (2009) N250 ERP Correlates of the Acquisition of Face Representations across Different Images. J Cogn Neurosci 21:625–641.

Kikuchi K, Masuda Y, Yamashita T, Kawai E, Hirao T (2015) Image analysis of skin color heterogeneity focusing on skin chromophores and the age-related changes in facial skin. Skin Research and Technology 21:175–183.

Kleiner M, Brainard D, Pelli D, Ingling A, Murray R, Broussard C (2007) What ’ s new in Psychtoolbox-3 ? Perception 36:1–16.

Kreibig SD (2010) Autonomic nervous system activity in emotion: A review. Biol Psychol 84:394–421.

Langeslag SJE, Gootjes L, van Strien JW (2018) The effect of mouth opening in emotional faces on subjective experience and the early posterior negativity amplitude. Brain Cogn 127:51–59.

Leutheuser H, Gabsteiger F, Hebenstreit F, Reis P, Lochmann M, Eskofier B (2013) Comparison of the AMICA and the InfoMax algorithm for the reduction of electromyogenic artifacts in EEG data. Proceedings of the Annual International Conference of the IEEE Engineering in Medicine and Biology Society, EMBS 6804–6807.

Lin L, Wang C, Mo J, Liu Y, Liu T, Jiang Y, Bai X, Wu X (2020) Differences in Behavioral Inhibitory Control in Response to Angry and Happy Emotions Among College Students With and Without Suicidal Ideation: An ERP Study. Front Psychol 11:543007.

MORETTI G, ELLIS RA, Mescon H (1959) Vascular Patterns in the Skin of the Face. Journal of Investigative Dermatology 33:103–112.

Nakajima K, Minami T, Nakauchi S (2017) Interaction between facial expression and color. Scientific Reports 2017 7:1 7:1–9.

Nakajima K, Minami T, Nakauchi S (2012) The face-selective N170 component is modulated by facial color. Neuropsychologia 50:2499–2505.

Nakajima K, Minami T, Tanabe HC, Sadato N, Nakauchi S (2014) Facial color processing in the face-selective regions: An fMRI study. Hum Brain Mapp 35:4958–4964.

Nguyen HN, Tamura H, Minami T, Nakauchi S (2023) The effect of facial colour on implicit facial expressions. Cogn Emot 1–8.

Palazova M, Mantwill K, Sommer W, Schacht A (2011) Are effects of emotion in single words non-lexical? Evidence from event-related brain potentials. Neuropsychologia 49:2766–2775.

Pelli DG (1997) The VideoToolbox software for visual psychophysics: Transforming numbers into movies. Spat Vis.

Peromaa T, Olkkonen M (2019) Red color facilitates the detection of facial anger — But how much? PLoS One 14:e0215610.

Pion-Tonachini L, Kreutz-Delgado K, Makeig S (2019) ICLabel: An automated electroencephalographic independent component classifier, dataset, and website. Neuroimage 198:181–197.

Polich J (2007) Updating P300: An integrative theory of P3a and P3b. Clinical Neurophysiology 118:2128–2148.

Quettier T, Maffei A, Gambarota F, Ferrari PF, Sessa P (2023) Testing EEG functional connectivity between sensorimotor and face processing visual regions in individuals with congenital facial palsy. Front Syst Neurosci 17:1123221.

Rossignol M, Philippot P, Douilliez C, Crommelinck M, Campanella S (2005) The perception of fearful and happy facial expression is modulated by anxiety: an event-related potential study. Neurosci Lett 377:115–120.

Rossion B, Caharel S (2011) ERP evidence for the speed of face categorization in the human brain: Disentangling the contribution of low-level visual cues from face perception. Vision Res 51:1297–1311.

Schindler S, Bublatzky F (2020) Attention and emotion: An integrative review of emotional face processing as a function of attention. Cortex 130:362–386.

Schupp HT, Junghöfer M, Öhman A, Weike AI, Stockburger J, Hamm AO (2004) The facilitated processing of threatening faces: an ERP analysis. Emotion 4:189–200.

Shibusawa M, Hasegawa Y, Tamura H, Nakauchi S, Minami T (2025) Dynamic versus static facial color changes: Evidence for terminal color dominance in expression recognition. J Vis 25:8–8.

Surcinelli P, Codispoti M, Montebarocci O, Rossi N, Baldaro B (2006) Facial emotion recognition in trait anxiety. J Anxiety Disord 20:110–117.

Tanaka JW, Sung A (2016) The “Eye Avoidance” Hypothesis of Autism Face Processing. J Autism Dev Disord 46:1538–1552.

Telzer EH, Mogg K, Bradley BP, Mai X, Ernst M, Pine DS, Monk CS (2008) Relationship between trait anxiety, prefrontal cortex, and attention bias to angry faces in children and adolescents. Biol Psychol 79:216–222.

Thorstenson CA, Elliot AJ, Pazda AD, Perrett DI, Xiao D (2018) Emotion-Color Associations in the Context of the Face. Emotion 18:1032–1042.

Thorstenson CA, McPhetres J, Pazda AD, Young SG (2022) The role of facial coloration in emotion disambiguation. Emotion 22:1604–1613.

Thorstenson CA, Pazda AD (2021) Facial coloration influences social approach-avoidance through social perception. Cogn Emot 35:970–985.

Thorstenson CA, Pazda AD, Krumhuber EG (2021) The influence of facial blushing and paling on emotion perception and memory. Motiv Emot 45:818–830.

Tsumura N, Haneishi H, Miyake Y (1999) Independent-component analysis of skin color image. JOSA A, Vol 16, Issue 9, pp 2169–2176 16:2169–2176.

Tsumura N, Ojima N, Sato K, Shiraishi M, Shimizu H, Nabeshima H, Akazaki S, Hori K, Miyake Y (2003) Image-based skin color and texture analysis/synthesis by extracting hemoglobin and melanin information in the skin. ACM SIGGRAPH 2003 Papers, SIGGRAPH ’03 770–779.

Willenbockel V, Sadr J, Fiset D, Horne GO, Gosselin F, Tanaka JW (2010) Controlling low-level image properties: The SHINE toolbox. Behav Res Methods 42:671–684.

Zonios G, Bykowski J, Kollias N (2001) Skin Melanin, Hemoglobin, and Light Scattering Properties can be Quantitatively Assessed In Vivo Using Diffuse Reflectance Spectroscopy. Journal of Investigative Dermatology 117:1452–1457.

